# Multivariate models of animal sex: breaking binaries leads to a better understanding of ecology and evolution

**DOI:** 10.1101/2023.01.26.525769

**Authors:** J. F. McLaughlin, Kinsey M. Brock, Isabella Gates, Anisha Pethkar, Marcus Piattoni, Alexis Rossi, Sara E. Lipshutz

## Abstract

‘Sex’ is often used to describe a suite of phenotypic and genotypic traits of an organism related to reproduction. However, these traits – gamete type, chromosomal inheritance, physiology, morphology, behavior, etc. – are not necessarily coupled, and the rhetorical collapse of variation into a single term elides much of the complexity inherent in sexual phenotypes. We argue that consideration of ‘sex’ as a constructed category operating at multiple biological levels opens up new avenues for inquiry in our study of biological variation. We apply this framework to three case studies that illustrate the diversity of sex variation, from decoupling sexual phenotypes to the evolutionary and ecological consequences of intrasexual polymorphisms. We argue that instead of assuming binary sex in these systems, some may be better categorized as multivariate and nonbinary. Finally, we conduct a meta-analysis of terms used to describe diversity in sexual phenotypes in the scientific literature to highlight how a multivariate model of sex can clarify, rather than cloud, studies of sexual diversity within and across species. We argue that such an expanded framework of ‘sex’ better equips us to understand evolutionary processes, and that as biologists it is incumbent upon us to push back against misunderstandings of the biology of sexual phenotypes that enact harm on marginalized communities.

## Introduction: Sex is not a single trait

In sexually reproducing species, the common assumption is that there are two sexes, strictly classified as female or male (Fisher 1930; Hurst 1996; Jaffe 1996). This assertion is supported by the cellular mechanisms of sexual reproduction, in which multiple (usually two) parental gametes combine genotypes to create an offspring with a novel genetic makeup. In animals, these gametes are of different sizes (e.g., large ova and small sperm), a condition called anisogamy. However, the binary classification of gametic sex breaks down when we consider the broader diversity of gametic phenotypes. For instance, hermaphroditic species possess both gamete types required for reproduction, and do not have separate sexes (Jarne and Auld 2006). Of the estimated 1.2 million animal species, roughly 5-6% are hermaphroditic (Jarne and Auld 2006). Outside of animals, systems of sex determination rely on genetic markers that determine compatibility between equally-sized gametes, a condition known as isogamy (Togashi and Cox 2011) found in fungi (Kothe 1996; Lee et al. 2010; Billiard et al. 2011, 2012), algae (Perrin 2012; Tillmann and Hoppenrath 2013), and amoebozoa (Douglas et al. 2016).

Operationally, the term ‘sex’ has two meanings - one, as a reproductive process that refers to the transmission of genetic information to the next generation, and another, as a categorical term that encompasses a broad collection of gametic, genetic, hormonal, anatomic, and behavioral traits (Gross 1996; Whitfield 2004; Engqvist and Taborsky 2016; Mank 2022). Whereas some biologists argue that gametes are the only meaningful sex categories (Goymann et al. 2023), we find several limitations to this gametic sex definition, particularly for ecologists and evolutionary biologists. First, gametes are rarely measured directly, with researchers instead relying on genetic and phenotypic proxies. Second, though biologists have drawn a direct connection between the evolution of gamete size and other sexual phenotypes (e.g., Kalmus 1932), these traits (gametes, genotype, hormones, anatomy, behavior) are not universally coupled. Joan Roughgarden summarized this problem: “the biggest error in biology today is uncritically assuming that the gamete size binary implies a corresponding binary in body type, behavior, and life history” (2013). Finally, selection typically acts on phenotypic traits, and variation in these traits is the raw material of evolution Thus, if we want to understand the evolution of diversity, we need to expand, rather than collapse, our definition of sex beyond binary categorization.

As a categorical term, ‘sex’ is often semantically flattened into a binary, univariate model, for which individuals are classified as either ‘female’ or ‘male’ (Figure 1A) in gonochoristic species (i.e., an individual produces one type of gamete throughout its lifespan). A more expansive definition of sex is bimodal - with most individuals falling within one of two peaks of a trait distribution (Figure 1B). However, even a bimodal, univariate model is an oversimplification, since ‘sex’ comprises multiple genetic and phenotypic traits, with variable distributions (see Figure 1C and case studies). Individuals may possess different combinations of chromosome type, gamete size, hormone level, morphology, and social roles, which do not always align or persist across an organism’s lifespan (Karkazis 2019; Griffiths 2021). Reliance on strict binary categories of sex fails to accurately capture the diverse and nuanced nature of sex.

**Figure 1:**
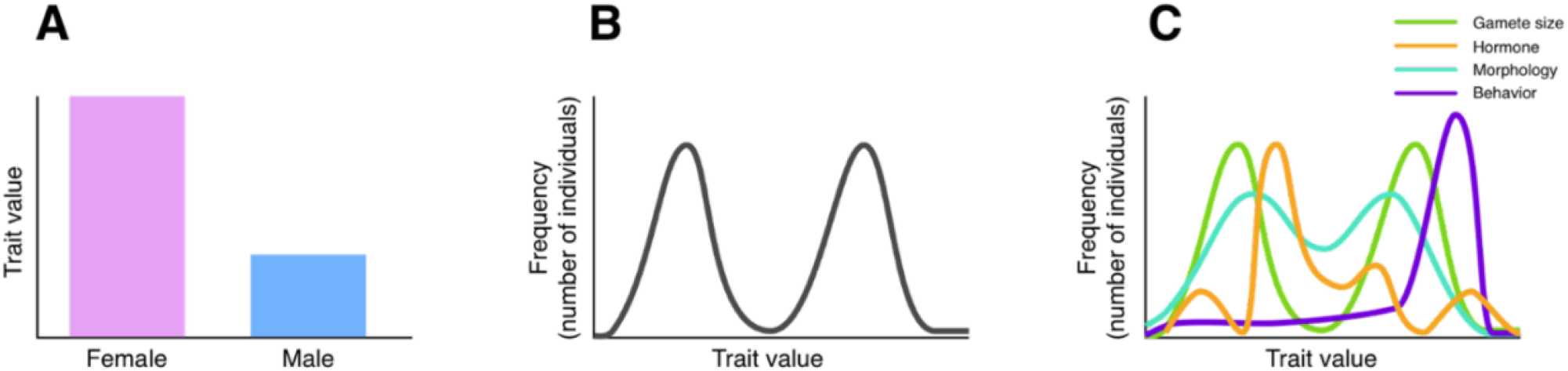
Three models of sexual phenotypes, demonstrated by their distribution across a hypothetical population: A) A strict binary, for which all individuals are unambiguously grouped into one of two categories. Whereas some traits, such as gamete size, operate this way, a binary does not accurately model the distribution of most phenotypic traits. B) A bimodal model, with most individuals falling around two peaks, or means, of a single sexual trait with continuous variation. C) A multivariate model of sex as a collection of traits, represented as individual lines, that contribute to the overall sex phenotype. Each trait has its own distribution, which may or may not be bimodal, or coincide with other traits.

Here we propose a model of animal sex as multivariate (Figure 1C), encompassing multiple traits that may or may not follow a binary distribution. Together, the distributions of these traits– gametic, hormonal, morphological, behavioral, etc.– comprise the sexual phenotype of an individual within a species. We lay out the shortcomings of assuming and analyzing sex as binary across these biological levels of organization. We then provide three case studies that illustrate the diversity of sex variation: decoupling sexual phenotypes in ‘sex-role reversed’ birds, the evolutionary consequences of a chromosomal inversion resulting in more than two sexes, and the ecological and evolutionary consequences of intrasexual color polymorphism in lizards. We argue that rethinking the default assumption of binary sex can help us better understand the evolutionary and ecological contexts in which these animals operate. We conduct a meta-analysis of terminology used to describe sexual phenotypes in scientific literature. Finally, we offer recommendations for future research and education on sex diversity.

We are certainly not the first, nor the last biologists to interrogate the definition of sex (Hoekstra 1990; Fausto-Sterling, 2012; Roughgarden 2013; Karkazis 2019), or the nature of biological categories more broadly (Cadena and Zapata 2021). We rely on literature mainly pertaining to organismal biology, spanning the fields of molecular genetics, neuroendocrinology, behavior, ecology, and evolution. We find it useful to clarify that our focus is primarily on animals, and vertebrates in particular. We also differentiate between the concepts of sex and gender (Goymann and Brumm 2018). Though the term ‘gender’ has occasionally been used to describe sexual phenotypes (Roughgarden 2013; Kutschera 2016), this term describes multifaceted and abstract self-interpretations of identity, an internal personal experience that expresses itself in the context of an individual’s specific culture, place, and time (Thorne et al. 2019). We validate the important role of gender in the human experience, especially in regard to transgender, nonbinary, and gender nonconforming (TNGC) identities. However, in studies of non-human animals it is key to differentiate between the specifically cultural role of gender and the observed traits of other organisms that we broadly call ‘sex’ (Goymann and Brumm 2018). Thus, throughout this paper we focus solely on sex, as anything similar to the human experience of gender in other organisms is fundamentally inaccessible to us.

### Sex at the genetic level

The genetics of sexual determination vary widely (Bachtrog et al. 2014; Figure 2). In the case of highly specialized sex chromosomes with reduced recombination, biologists classify individuals as homogametic (i.e., have two of the same sex chromosome type) or heterogametic (having two distinct chromosome types), with sexual phenotypes following from those genotypes. However, even these ‘simple’ systems include intermediate (intersex) phenotypes (Vaiman and Pailhoux 2000), as additional genes outside the sex chromosomes modulate developmental pathways. The most widely familiar systems are XX/XY, where the presence of certain sex-determining genes, such as *sry* on the Y chromosome in most mammals, activates diverging pathways early in the developmental process (Bachtrog et al. 2014; Stévant et al. 2018) (Figure 2A). ZZ/ZW systems operate similarly, but with the ova-producing individuals being heterogametic (Bachtrog et al. 2014) (Figure 2B). In some taxa, rather than a small number of genes on the smaller chromosome, it is the dose-dependent effects of homogametic chromosomes that lead to higher expression of genes that start a particular development pathway (Smith et al. 2009). Dose-dependent effects also underlie the haplodiploid systems of many insects, where unfertilized and fertilized ova develop into different social and reproductive classes (Beye et al. 2003; Verhulst et al. 2010; Figure 2C). Some combination of sex-determining genes and dosage dependence may be at play in species for which the smaller sex chromosome and most of its associated genes have been lost entirely (Sutou et al. 2001; Just et al. 2002; Kuroiwa et al. 2010).

**Figure 2:**
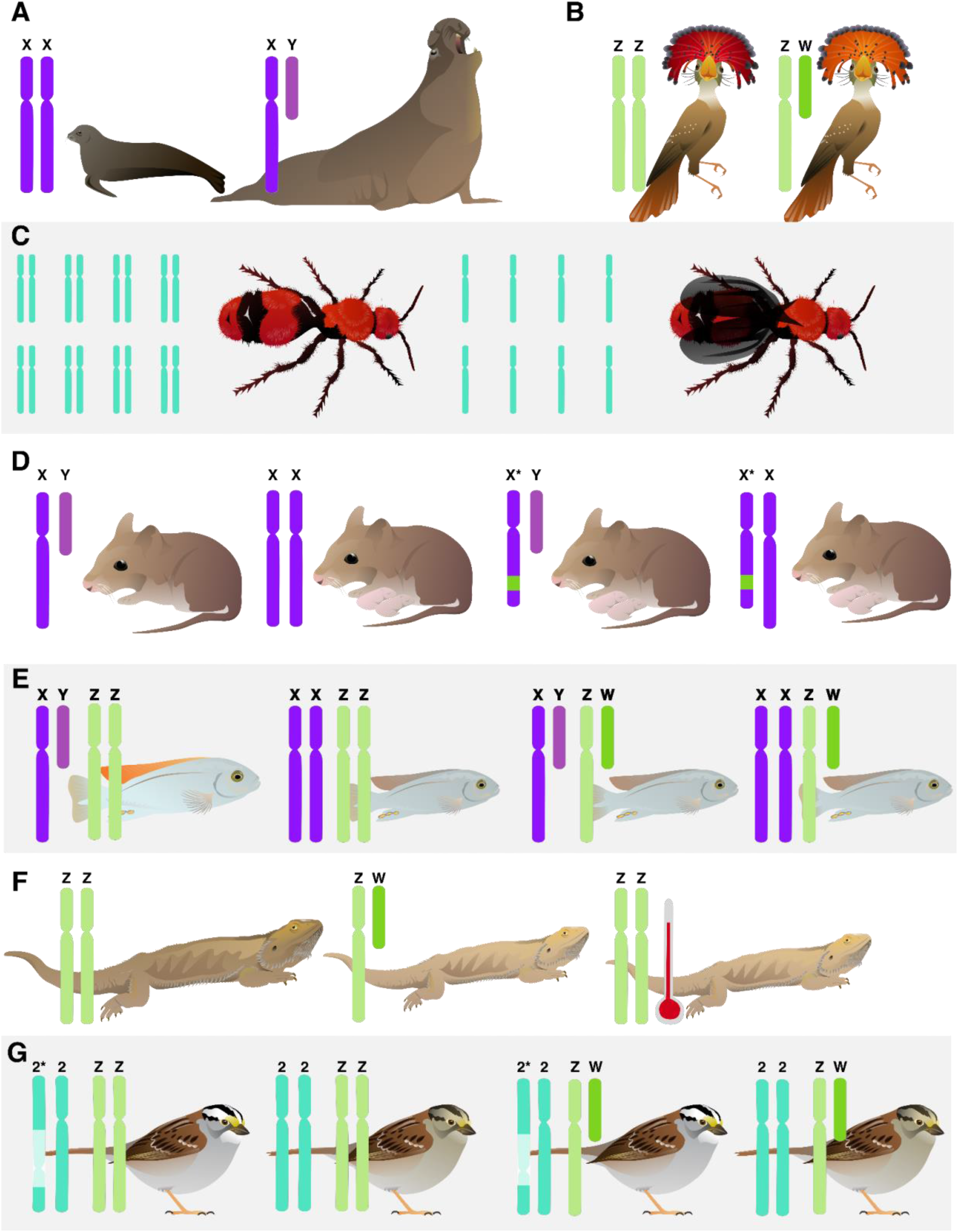
Mechanisms of sex determination vary widely across animals. The most familiar (top row) are the relatively simple (**A**) XX/XY (in purple, here shown with northern elephant seal, *Mirounga angustirostris*, as an example taxon) and (**B**) ZZ/ZW (in green, shown with royal flycatcher, *Onychorhynchus coronatus*). (**C**) Haplodiploid sex determination involves sex being determined by having one or two copies of the genome, and is found predominantly in insects such as the pictured *Dasymutilla occidentalis*. (**D**) A mutation in the X chromosome of some African pygmy mice (*Mus minutoides*) overrides the presence of a Y chromosome, and (**E**) in some cichlids (shown here with *Metriaclima mbenjii*), both the absence of the W chromosome and presence of the Y are required for an individual to develop ‘male’ characteristics. (**F**) In central bearded dragons (*Pogona vitticeps*), the ZZ genotype is overridden by high temperatures, leading to phenotypically ‘female’ individuals capable of laying eggs. (**G**) The white-throated sparrow (*Zonotrichia albicollis*) is found in two color morphs linked to a large inversion on chromosome two (in turquoise), which is paired with the Z and W genotype to determine which individuals typically reproduce with each other.

In polygenic systems, multiple loci on several chromosomes, both sex-linked and autosomal, interact to determine sexual phenotypes. Polymorphisms in the sex chromosomes can override sex-determining genes (Figure 2D; Veyrunes et al. 2010; Saunders et al. 2014; Zhao et al. 2017) or lead to multiple reproductive morphs (Sandkam et al. 2021). In several species of cichlid fish, sex is determined by the interplay of both ZZ/ZW and XX/XY chromosomes, resulting in four genotypes but two phenotypes (Ser et al. 2010; Moore et al. 2022; Figure 2E). In species with these polygenic systems, the number of reproductive phenotypes is not necessarily the same as the number of genotypes in the population, demonstrating how genotype alone is not sufficient in defining sexual categories.

Not all sexual differentiation is controlled by genetic mechanisms, in part or whole (Bachtrog et al. 2014). Temperature-dependent sex determination is common among reptiles (Figure 2F) (Sarre et al. 2004) and fish (Baroiller et al. 2009). In its simplest form, the temperature of incubation determines which hormonal pathways are triggered in a developing egg (Sarre et al. 2004). In other systems, temperature can override genetic pathways, resulting in animals with the genetic markers usually associated with one sex, but the phenotypic traits of another (Quinn et al. 2009; Holleley et al. 2015; Whiteley et al. 2017; Wiggins et al. 2020). Other environmental variables may likewise interact with genetic sex determination, possibly through epigenetic factors (Cabej 2018; Piferrer 2021), but these last mechanisms are only just beginning to be explored. Altogether, evidence from these diverse species reveals that genetic sex is not the universal binary often assumed, and that genetic sex is one variable among the many traits that comprise multivariate sex.

### Sex at the endocrine level

Hormones influence a number of sexual phenotypes, including 1) morphological traits such as gonads, genitals, ornamentation, and body size, 2) reproductive states such as estrous cycling and gametogenesis, and 3) behaviors such as courtship displays, receptivity to mating, aggression, and parental care (Adkins-Regan 2005). So-called ‘sex’ hormones typically refer to steroid hormones produced in the testes and ovaries, and include estrogens, progestogens, as well as several androgens (testosterone, androstenedione, dihydrotestosterone, and 11-ketotestosterone). However, many of these hormones are also produced outside of the gonads, including in the adrenal cortex, fat, and brain (Schmidt et al. 2008, Saldanha et al. 2011). Furthermore, variation in other elements of endocrine signaling pathways (e.g., receptors, enzymes, steroid binding proteins) beyond circulating hormone levels may contribute to the expression of sexual phenotypes (Adkins-Regan 2005, Hau 2005). For instance, in male zebra finches (*Taeniopygia guttata*), testosterone is converted to 17β-estradiol via the enzyme aromatase in the brain, which then regulates directed singing (Walters et al. 1991). In another songbird, dark-eyed juncos (*Junco hyemalis*), variation in the expression of genes encoding the androgen receptor, estrogen receptor, and aromatase enzyme correlates with aggression in both sexes (Rosvall et al. 2012). Thus, androgens and estrogens are not strictly associated with regulating the expression of ‘male typical’ and ‘female typical’ morphologies and behaviors. However, a common misconception is that androgens regulate ‘masculine’ phenotypes and estrogens regulate ‘feminine’ phenotypes, which has led to the widely used terms ‘masculinization’ and ‘feminization’. We argue that this terminology is circular and confusing. Steroid hormones are not sex-specific - both females and males possess the capacity to produce, metabolize, and respond to these hormones (Staub and De Beer 1997). Thus, testosterone is not a ‘male hormone,’ and estrogen is not a ‘female hormone’.

Whereas it is often the case that average peak hormone levels may be higher or lower in one sex, this depends on various contexts, including when animals are developing (organizational effects) and when they are breeding (activational effects; Phoenix et al. 1959), as well as in response to social stimuli (Wingfield et al. 1990; Goymann et al. 2019). Reflecting back on our multivariate framework, testosterone levels in circulation are typically higher in adult breeding males than in females for many species (Goymann and Wingfield 2014; Van Leeuwen and Bladh 2016), reflecting the bimodal distribution depicted in Figure 1B. However, there are many hormonal patterns that follow the distributions depicted in Figure 1C. For example, testosterone in many temperate bird species peaks seasonally during territorial establishment and mating, but decreases to basal, non-detectable levels during the non-breeding season, when gonads regress (Wingfield et al. 2019). Within an individual, testosterone levels vary substantially throughout the day (Panico et al. 1990; Brambilla et al. 2009; Greives et al. 2021). Given their daily, seasonal, and cross-species variation, hormone levels are not a binary indicator of sex.

### Sex at the morphological level

Organismal biologists often assign binary sex to species based on their primary sexual characteristics, despite the diversity of genital morphology within and among taxa. For example, spotted hyenas (*Crocuta crocuta*) mate, give birth, and urinate through a urogenital canal at the tip of a hypertrophied clitorophallus (Glickman et al. 2006), which they may use to signal social status (Hofer and East 1995) or to facilitate post-copulatory sperm choice (Holekamp 2006). In *Neotrogla* barkflies, females have a highly elaborate penis (gynosome) that retrieves sperm from the male’s vagina (Yoshizawa et al. 2014). In color polymorphic lizards, male color morphs have distinct hemipene morphology associated with their alternative mating strategies (Gilman et al. 2019). Phallus polymorphism is common in hermaphroditic gastropods, in which some individuals have a functional penis (euphally), some have a non-functional penis (hemiphally), and some have no penis (aphally) (Leonard et al. 2007). In birds, only 3% of species even have phalluses (external genitalia) (Brennan 2022). As with phallic diversity, animal morphologies involved in birthing and brooding exhibit extraordinary intra- and intersexual variation. Male pregnancy has evolved several times in syngnathid fishes (seahorses and pipefish) where gestation in males requires morphological and physiological changes in paternal tissues to incubate developing embryos in a specialized brooding pouch, analogous to structures in viviparous females (Carcupino et al. 2002). In many birds, for which biparental care is common (Clutton-Brock 2019), any adult can develop brood patches to incubate eggs (Tucker 1943). In mammals (humans included), genital structures are not formed de novo in males and females, but develop from a shared anatomical basis that can differentiate into forms other than the two most commonly observed pathways, leading to intermediate phenotypes that are unfortunately pathologized (Quigley et al. 1995; Dreger 2006; Grimstad et al. 2021). Thus, as a result of variation in developmental pathways, external genitalia are not necessarily concordant with sex chromosome makeup, and the presence, shape, and size of genitalia do not always fall into binary categories of morphological sex.

Secondary sexual characteristics include ornamentation, color, weaponry, and many other sexual morphologies that are not directly involved in reproduction. Although these traits are expected to be sexually dimorphic, there are many exceptions. For example, polymorphisms, both sex-limited and not, present a classic example of more than two sexual phenotypes (Mank 2022). The three male morphs of the ruff (*Philomachus pugnax*), a bird in the sandpiper family, exhibit extremely divergent size and plumage ornamentation, which correspond to alternative mating strategies and chromosomal genotypes (Küpper et al. 2016). In many odonates there are multiple female morphs, consisting of one or more ‘gynomorphs’ that vary in color, and one ‘andromorph’ that closely resembles males in color, sexual ornamentation, and behavior, presumably to avoid mating harassment (Gallesi et al. 2015). These so-called ‘andromorphs’ exhibit morphological and behavioral variation distinct from males and ‘gymnomorph’ females (Paulson 1998; Fincke et al. 2005; Gallesi et al. 2015), resulting in an overall phenotype that does not conform to a binary distribution and likely arises due to a variety of factors beyond male mimicry (Fincke et al. 2005). For both primary and secondary sexual morphologies, there are many cases in which the distribution of these traits is continuous, rather than binary.

### Sex at the behavioral level

Scientific notions of how animals ‘should’ adaptively behave originate with Darwin in the Victorian era (Darwin 1871). The Darwin-Bateman paradigm posits that ‘traditional’ sex roles arise from anisogamy, such that mate competition is stronger in the sex with the smaller gametes (Bateman 1948; Dewsbury 2005). Trivers (1972) extended this paradigm with the hypothesis that gametic investment drives postzygotic parental investment, which should be higher in females because of larger gamete size. Theories on the evolution of sex roles have dominated sexual selection research, but not without criticism (Gowaty et al. 2012; Ah-King 2013; Tang-Martínez 2016), as others have argued for the importance of ecology, sex ratio, and life history traits on the evolution of sex-specific behaviors (Kokko and Jennions 2008; Mokos et al. 2021; Kappeler et al. 2022). Indeed, there are many examples of competitive females and choosy males across animal groups (reviewed in Hare and Simmons 2018; Edward and Chapman 2011). Outside of mammals, many examples in fish, frogs, and birds contradict the notion that the sex with the larger gamete suffers higher mating costs and invests more resources into the next generation, such as the repeated evolution of male parental care in frogs (Townsend et al. 1984; Gross 2005; Furness and Capellini 2019).

Within species, alternative reproductive strategies are a classic example of discontinuous variation in behavior (Gross 1996; Sinervo and Lively 1996; Oliveira et al. 2008; Stiver et al. 2015), and can arise via different mechanisms of genotype, condition dependence, and social environment. For example, in the bluegill sunfish (*Lepomis macrochirus*), there are three different reproductive strategies found in males, one of which overlaps with courtship soliciting behaviors in females (Dominey 1980). The diversity of sexual behaviors also includes the common occurrence of same-sex sexual behavior (SSB) in animals (Monk et al. 2019). SSBs have been observed across a large number of animals (Bailey and Zuk 2009), but were frequently unreported by early naturalists due to societal biases (Russell et al. 2012).

Compared to the other traits we review, sex studied at the social level seems to more readily acknowledge variation outside of binary categories, perhaps because plasticity is a fundamental aspect of behavioral research. Furthermore, several recent studies in animal behavior and evolutionary biology have acknowledged that the positionality of the scientists themselves influences their research (Ahnesjö et al. 2020; Tang-Martínez 2020; Ah-King 2022; Pollo and Kasumovic 2022), making more space for scientists to reflect on how their identities shape their scientific practices.

We next provide three case studies, which highlight how a de-constructed approach to sex can lead to asking novel empirical questions. These examples come from our own study systems and research expertise. Rather than considering these systems as rare exceptions, we can use them as models to understand the diversity of sexual phenotypes across multiple biological levels.

### Case study 1: ‘Sex-role reversal’ and the decoupling of sexual phenotypes

Binary expectations of sexual phenotypes are often met with exceptions, and one classic example are birds with polyandrous mating systems, including black coucals (*Centropus grillii*), spotted sandpipers (*Actitis macularius*), buttonquails (Turnicidae), and northern jacanas (*Jacana spinosa)*. In these systems, individuals with larger gametes typically face stronger sexual selection for mating opportunities (Fritzsche et al. 2021), decoupling sexual phenotypes from anisogamous expectations. These systems are often called ‘sex-role reversed’ because they defy ‘traditional’ expectations of sex roles historically found in mammals - male competition and female parental care. However, sex role terminology imposes binary categorization, limiting our understanding of continuous variation in phenotypes (Ah-King and Ahnesjö 2013). As we describe below, polyandrous mating systems are not simply the ‘reverse’ of polygynous mating systems, but rather, they are novel evolutionary outcomes to be studied in their own right, not solely in relation to a polygynous foil. We propose that studying so-called ‘exceptional’ systems can help reveal unexamined routes of phenotypic evolution, expanding the conceptual framework for how we study sexual diversity.

Here, we review examples from northern jacanas (*Jacana spinosa*), a socially polyandrous shorebird and a focal species of one of our co-authors. Jacanas allow us to evaluate: To what extent do the sexes differ in the first place? For which traits, and at what developmental and seasonal timepoints? At the genetic level, jacanas have ZW/ZZ sex determination and females are heterogametic - this categorization, as for most birds, is binary. At the endocrine level, circulating testosterone concentrations are context-dependent. Females have significantly lower testosterone than males in the courtship stage, but sex differences in testosterone disappear when males are conducting parental care (Lipshutz and Rosvall 2020a). Thus, the degree of sexual dimorphism in testosterone varies with male breeding stage, a pattern that applies broadly across independent origins of social polyandry (Eens and Pinxten 2000; Lipshutz and Rosvall 2020b). The gonadal size of territorial males also fluctuates between reproductive stages of courtship and parental care, whereas territorial females remain in a continuous state of fertility (Lipshutz and Rosvall 2020a). For this species, the cyclical variation in hormonal and gonadal traits reveals that these sexual phenotypes, and their degree of sexual dimorphism, are dynamic.

At the morphological level, sexual size dimorphism is biased towards females, who have more exaggerated ornamentation, weaponry, and body size (Emlen and Wrege 2004) There is some evidence that wing spur length positively correlates with testosterone in females, but not males (Lipshutz and Rosvall 2020a), suggesting that hormonal sensitivity may vary between the sexes. At the social level, female jacanas defend territories that encompass multiple male mates (i.e. social polyandry), and males conduct the majority of parental care (Emlen and Oring 1977). Despite expectations that ‘role-reversed’ male jacanas should be choosy, virtually nothing is known about mate choice in this system. Thus far, competition for territories seems to be the predominant determinant of mating success in jacanas (Emlen and Wrege 2004). Larger females and males gain access to breeding territories, whereas floater females and males do not reproduce. Thus, reproductive access varies within and among both sexes, and there are non-reproductive phenotypes for both females and males. Altogether, the genetic, endocrine, morphological, and social traits of ‘sex-role reversed’ species such as northern jacanas demonstrate that sexual phenotypes are dynamic, complex, and support a multivariate framework of sexual diversity (Figure 1C, 3).

**Figure 3:**
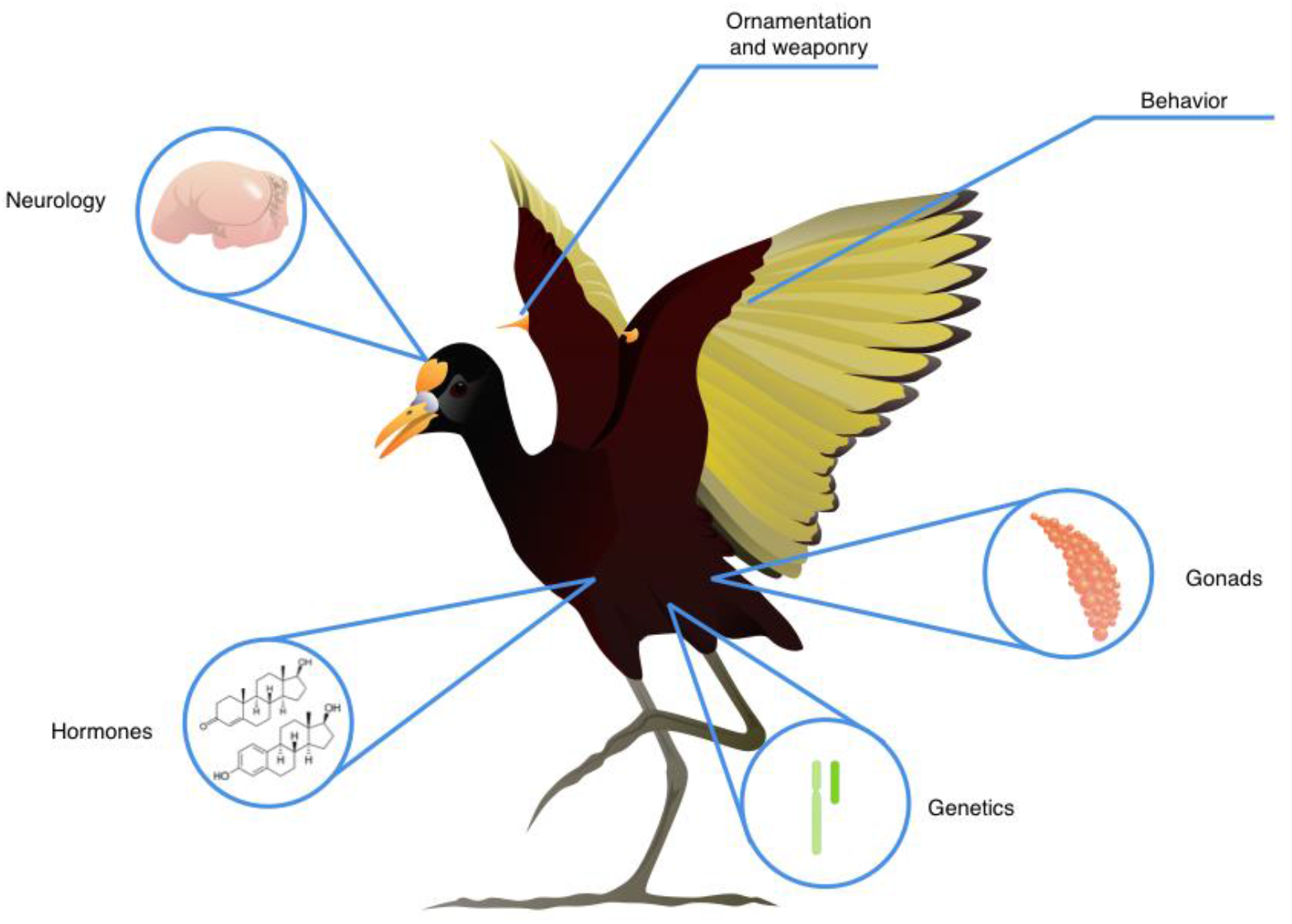
Multivariate sex in the northern jacana. ‘Sex’ includes phenotypic variation at multiple levels, from genetic makeup, anatomy and morphology, hormone levels, and social roles and behaviors.

### Case study 2: The evolutionary consequences of more than two sexes

How does the number of operative sexes influence the evolution of a species? White-throated sparrows (*Zonotrichia albicollis*), have been described as ‘the bird with four sexes’, with both tan and white stripe morphs occurring in ZZ and ZW individuals. This species mates disassortatively by color morph, with tan stripe morphs predominantly mating with white stripe morphs (Hedrick et al. 2018). The more aggressive white-stripe morph has a large inversion on chromosome 2 that functions as a ‘supergene’, which tan-stripe individuals lack (Tuttle et al. 2016; Falls and Kopachena 2020; Maney et al. 2020;Figure 2G). These morphs are further associated with differences in testosterone and estrogen (Maney and Küpper 2022). Population persistence as a whole requires that all four morphs be present (Campagna 2016; Maney et al. 2020). A similar system occurs in hybrid *Pogonomyrmex* ants, where each colony requires three sexes to operate, and four to persist beyond a single generation (Parker 2004).

There are significant evolutionary consequences to having more than two operative sexes. A key parameter in population genetics is effective population size (*N*_*e*_; Charlesworth 2009), which is fundamentally impacted by the number of sexes in a given population (Caballero 1994). Assuming an equal sex ratio, the absolute maximum of *N*_*e*_ will be equal to 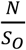, where *N* is the census population size and *S*_*o*_ is the operative number of sexes. As *N*_*e*_ influences the strength of selection, genetic drift, and genetic diversity of a population (Charlesworth 2009), the operative number of sexes substantially impacts the evolutionary trajectory of a lineage. The number of sexes in a system also shapes genome evolution (Charlesworth and Mank 2010), mating kinetics (Power 1976; Iwasa and Sasaki 1987), and sexual selection (Emlen and Oring 1977), and thus a framework of sex that accounts for diversity in sexual phenotypes allows us to more accurately model evolutionary processes (Mokos et al. 2021).

We can gain novel insights by applying this framework to other taxa with genetic polymorphisms. The previously mentioned ruff also has a genetic polymorphism linked to multivariate phenotypic differences in reproductive strategies (Lank et al. 1995; Jukema and Piersma 2006). It is typical to frame such systems as having *subtypes* of two sexes– in this case, three males and one female. But if we are willing to consider the white-throated sparrows and *Pogonomyrmex* systems as having more than two sexes, why not apply a similar framework to the ruff and other polymorphic systems (Galeotti et al. 2003; Roulin 2004)? Our binary framing of sex in these systems may interfere with understanding their evolutionary trajectories.

### Case study 3: Ecological and evolutionary consequences of multimodal sex in color polymorphic lizards

Color polymorphism, the evolution of two or more genetically-determined color morphs within a single population (Ford 1945; Sinervo et al. 2001; Andrade et al. 2019), is a classic example of multivariate and multimodal sexual diversity (Figure 1C; Mank 2022). Intrasexual color morphs have evolved in every major animal group across the tree of life, and are particularly common in lizards (Cote et al. 2008; Stuart-Fox et al. 2021; Brock, McTavish, et al. 2022a). In lizards, intrasexual color morphs occur in both females and males, and are often associated with alternative sexual phenotypes comprising discontinuous variation in genital morphology (Gilman et al. 2019), clutch size (Sinervo et al. 2000), mate preference (Pérez i de Lanuza et al. 2013), mating strategies (Gross 1982; Sinervo and Lively 1996), and endocrine profiles (Huyghe et al. 2009; Brock et al. 2020). These distinct sexual phenotypes give rise to ecological specialization among morphs, including thermal preference (Thompson et al. 2023), microhabitat selection (BeVier et al. 2022), diet (Lattanzio and Miles 2016), and non-sexual behaviors (Brock and Madden 2022; Brock, Chelini, et al. 2022b). Although each morph utilizes a specialized set of resources, in aggregate, greater morph diversity can contribute to a greater niche breadth in polymorphic populations relative to monomorphic populations (Pérez i de Lanuza et al. 2018; Forsman et al. 2008); Figure 4). For example, in *Podarcis* wall lizards with three genetically-determined throat color morphs (Figure 4), morphs have different preferred body temperatures and microhabitat use that results in morphs that use a narrow space within the overall species’ environmental niche space (Pérez i de Lanuza and Carretero 2018; Thompson et al. 2023). As an ecological consequence, the overall greater niche breadth of polymorphic populations is hypothesized to confer greater capacity for range expansion, less susceptibility to environmental change, and a larger buffer to extinction risk (Forsman et al. 2008). Evolutionary consequences of multimodal morphs include punctuated patterns of trait evolution, greater mobility on the adaptive landscape, and increased diversification rates (West-Eberhard 1986; Corl et al. 2010; Hugall and Stuart-Fox 2012; Brock, McTavish, et al. 2022a). New species could form from different morphs if they become reproductively isolated or fixed within a population (West-Eberhard 1986; Gray and McKinnon 2007); (Seehausen et al. 2008; Corl et al. 2010; McLean and Stuart-Fox 2014; Brock, Madden, et al. 2022c). In addition to facilitating speciation, character release associated with morph fixation could produce punctuated accelerations of morphological change (Eldredge 1976; West-Eberhard 1986; Corl et al. 2010). Thus, the ecological differences that emerge from intrasexual polymorphism are a crucial component of biodiversity within a species, which has far-reaching consequences for population persistence in a rapidly changing world. Collapsing intrasexual polymorphisms into a female-male binary erases extensive multivariate phenotypic variation (Figure 4), including its ecological and evolutionary consequences.

**Figure 4:**
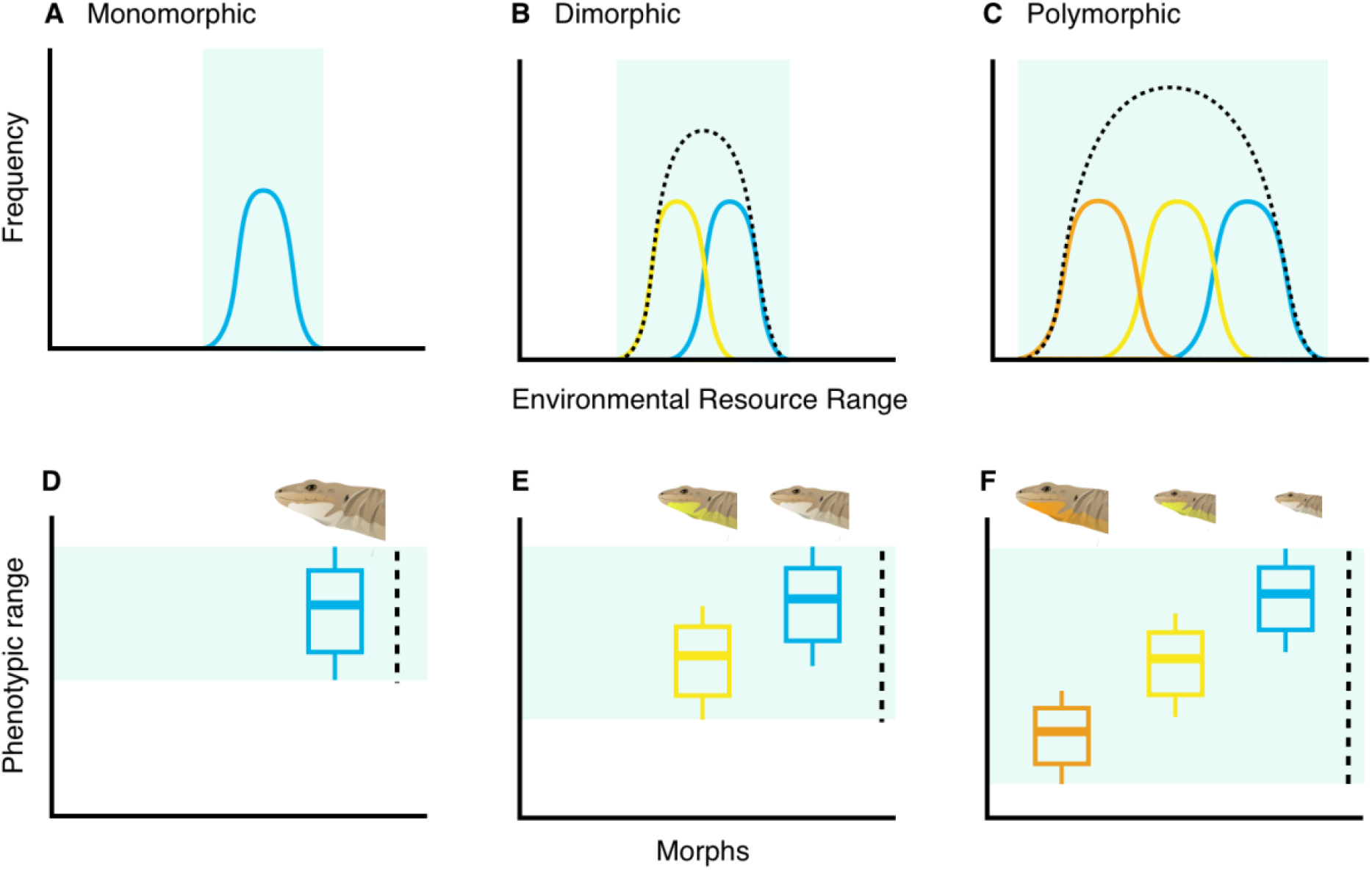
Illustration of alternative patterns of population-level environmental niche breadth (dotted line) distributions for A) monomorphic, B) dimorphic, and C) polymorphic populations of wall lizards (*Podarcis*). Different color morphs (solid color lines) have non-overlapping environmental resource use that, in the aggregate, result in an overall greater niche breadth in an A) monomorphic population compared to B) dimorphic and C) polymorphic populations.

These case studies demonstrate that breaking the sex binary - and expanding to a multivariate model of sex - is a useful framework for understanding animal ecology and evolution. Each of these animal systems possesses a different set of genetic, endocrine, morphological, and behavioral sexual traits, which do not fit neatly into binary distributions. Once we detach our assumptions of binary sex classifications, we can ask more expansive questions such as ‘how does variation within and among these sexual traits shape ecological and evolutionary processes?’

### Literature survey: How does terminology for sexual diversity change over time?

The language used in published research forms the foundation of how we as scientists think about, discuss, and investigate the natural world. To track how terminology usage has changed over time in ecology and evolutionary biology, we conducted a systematic review on Web of Science of search terms associated with genetic, endocrine, morphological, and behavioral levels of sex. Our search terms included the following: “andromorph/gynomorph” (60 papers), “female/male hormone” (40) “female-like/male-like” (124), “feminize/masculinize” (539) as well as outdated and transphobic language, including “transsexual”, “transvestite” and “she-male” (7). We also surveyed the conflation of sex and gender (1047). Additional details are available in the Supplementary Materials.

We initially identified 2,222 papers, 1,817 of which were exclusive to non-human animals, could be assigned a taxonomic class, and were available to Loyola University Chicago online (Table S1, Supplementary Materials). We used a chi-square test to evaluate taxonomic bias associated with each term, and a line of best fit to examine change in search term usage over time.

We found that most terms were significantly associated with a taxonomic class, except for masculinize/feminize and gender/sex conflation (Table S2). Andromorph/gynomorph are almost exclusively used in insects, particularly damselflies. “Female-like” and “male-like” are most commonly used in birds. Application of the outdated and offensive term “she-male” was confined to reptiles, garter snakes (*Thamnophis* sp.) in particular. Most terms in our literature survey increased in frequency over time, except for “she-male” (Figure S2).The conflation of sex and gender increased in the past three decades (Figure 5b), consistent with findings from a previous study (Goymann and Brumm 2018).

**Figure 5.**
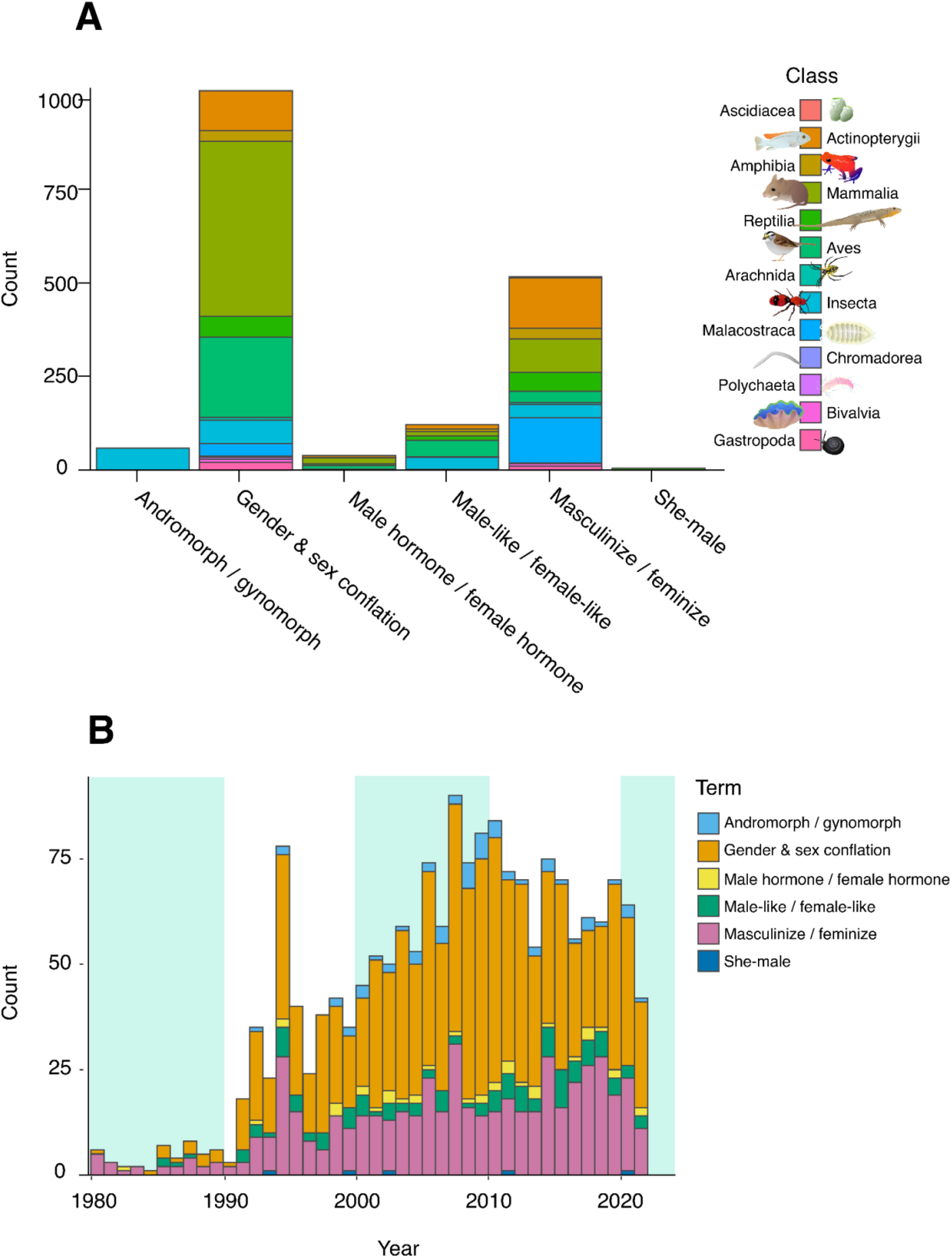
Literature survey for terms (a) by taxonomy and (b) over time. The search was conducted on 18 October 2022.

Terminology describing sex can have different meanings for different groups of people. Language is influenced by culture, and this can shift rapidly over time. What is popular today may not necessarily be seen as appropriate in a few years. Thus, it is important for biologists to be explicit in how we use these terms in our work. Sex is complex, so talking about it requires context and specificity - are we inferring sexes genetically? From their gonads? By color? Taking this path forward is important to not only reduce harm and improve science, but also to encourage and facilitate more research in systems and species that do not fit our own societal perceptions of what is “normal” when it comes to sexual phenotypes.

### The interplay of science and society

Multivariate models of sex reveal overlapping but not necessarily coincident phenotypes at every biological level within an individual, from the molecular to the behavioral (Maney, 2016). In zoology, we impose a binary categorization of sex as an emergent property of many traits. Whereas some of these traits do typically have a bimodal distribution (some chromosomes, gametes), others demonstrate largely continuous or multimodal variation (hormone levels [(Wingfield et al. 1990), morphology [Mank 2022], behavior [Dominey 1980)]), suggesting that most animals can best be studied from the framework of multiple phenotypic axes– some categorical, but most continuous. Even the basic inclusion of sex as a variable is missing from many studies, particularly in fields related to human health (Woitowich et al. 2020; Garcia-Sifuentes and Maney 2021). However, uncritically applying a simple binary without considering the mechanisms shaping sex-specific effects can confound inferences (Casto et al. 2022) and when applied to humans, completely erases the biological realities of trans, nonbinary, and gender-nonconforming (TGNC) and intersex people (Cheung et al. 2021; Phiri-Ramongane and Khine 2022).

Though we focus on animals, embracing complex models of sex has also led to greater understanding of fungi and plants (Lee et al. 2010; Billiard et al. 2011, 2012). For example, exploration of the diversity of fungal sex has illuminated the evolution of the process of sexual reproduction (Lee et al. 2010), the advantages and disadvantages of anisogamy (Billiard et al. 2011), and the evolutionary success of fungi in general (Naranjo-Ortiz and Gabaldón 2020). Here, we have argued that applying such a lens to animals may lead to similarly novel insights. Some readers may consider our examples of sexual diversity as rare anomalies – exceptions to the evolutionary rule, which do not require a novel framework to explain. We acknowledge some variations in sexual phenotypes may be relatively less common in animals, but that does not minimize their importance in our understanding of evolution. When we regard diversity as a deviation from the norm, we ignore its successful role in generating novel solutions to evolutionary challenges. Our frameworks of sex need to accommodate and embrace this diversity, lest we fall prey to a deterministic model of evolution, where the ideal endpoint is always binary.

As biologists, we seek to understand the vast diversity of life in all its wonderful forms - yet the lens we use fundamentally shapes that which we can observe. The historical legacies of sexism, racism, queerphobia, and ableism have deeply influenced the frameworks we use to study nature (Branch et al. 2022; Graves et al. 2022; Kamath et al. 2022), and the exclusion of certain groups of people from the scientific enterprise has delayed important discoveries such as the prevalence of female bird song (Haines et al. 2020) and the existence of hemiclitores in snakes (Folwell et al 2022a; Folwell et al 2022b). Challenging these foundations is difficult but vital to both increasing inclusion in biology (Hales 2020; Casper et al. 2022) and dismantling assumptions that interfere with our ability to observe and interpret the natural world (Monk et al. 2019; Ahnesjö et al. 2020; Kamath et al. 2022; Packer and Lambert 2022).

We recognize that TGNC and intersex people are valid regardless of the sexual diversity we find in nature. Human rights cannot be defined solely by biology - this is a prime example of the appeal to nature fallacy. However, opposition to the inclusion of TGNC and intersex people in society is often based in the language of biology. Especially in the United States, legislation targeting TGNC people is increasingly undergirded with the simplistic binary model of ‘biological’ sex (such as OH HB454 §3129.02 2021, WV HB3293 §18-2-25d.b1 2021, MO SB22 §191.1720 2022, and TX HB672 88R §71.004.1A 2022). There is pressure for scientists to avoid making the politics of our work explicit, especially those of us who do not directly study social issues. However, our science is being weaponized to discriminate against marginalized groups (Ha et al. 2014). It is imperative that we biologists challenge the misuse and abuse of this language (Miyagi et al. 2021) and confront how our scientific models impact society (Bazzul and Sykes 2011). As biologists, it is our responsibility to dispel misconceptions and recognize the rich tapestry of diversity in nature.

## Acknowledgements

Thanks to Kelsey Lewis and Sam Sharpe for the opportunity to contribute to this symposium. Aqdas Aftab, Wolfgang Goymann, and Kim Rosvall provided fruitful conversations discussing sex and gender diversity. Claire Curry and Elliott Smith provided vital assistance in designing the systematic review and mapping analysis. Thank you to Savvy Cornett in the Hofmann Lab at UT Austin and Kwasi Wrensford in the UC Berkeley Animal Behavior Reading Group for invitations to discuss the preprint version of this paper. Anusha Bishop, Anne Chambers, Jay Falk, Ambika Kamath, Daniel Olivera, Erin Westeen, Alejandro Rico-Guevara and four anonymous reviewers provided valuable feedback on the manuscript. Thank you to the Trans Formations Project, whose database of legislation was very helpful in understanding the use of the term ‘biological sex’ in recent legislation – we encourage interested readers to follow their tracking efforts at https://www.transformationsproject.org/.

