## Supplementary Materials for "Multivariate models of animal sex: breaking binaries leads to a better understanding of ecology and evolution"

**Supplementary Material**

***Literature survey methods***

***We completed our search on 18 October, 2022.*** We searched for literature in the Web of Science Core Collections, limited to English-language papers in journals subscribed to by Loyola University Chicago: Science Citation Index Expanded (1980 - present), Social Sciences Citation Index (1980 - present), Arts & Humanities Citation Index (1980 - present), and Emerging Sources Citation Index (2017 - present). Abstracts or posters from conferences were not included.

We designed our search strings to exclude humans, plants, fungi, and bacteria (Table 1). We further refined our search to the Zoology Research Area. For each result, the study organism(s) were recorded and assigned to a taxonomic Class. We excluded studies that 1) covered taxonomic groups belonging to more than one Class, 2) could not be assigned to a Class based on unclear taxonomy, 3) were not about animals, and 4) we could not access via Loyola University Chicago’s library system. We defined the remaining studies as “usable papers” (Table S1).

For gender/sex, we determined whether the terms were conflated by evaluating whether the terms were used interchangeably, or whether they were defined as separate. We only included cases where gender and sex were used interchangeably. The terms “transsexual” and “transvestite” were only found once each, in Reptilia and Insecta respectively, and thus not included in the summary figures (Figure 5a, S1).

To evaluate potential taxonomic bias for each search term, we conducted a chi-square test. We compared the percentages of each taxonomic class in a specific search term, relative to the percentage of each class in our total literature survey. Although there may be a taxonomic bias in the literature for which classes of animals are better studied than others (e.g. mammals vs. bivalves), we did not account for this bias. Nor did we account for the relative differences in taxonomic diversity within one class relative to another (e.g. insects vs. amphibians).

To examine how search term use has changed over time, we produced lines of best fit for each search term. We did not correct for the general increase in the number of papers published over time.

Table S1. Literature survey search terms, search string specifications, number of initial papers recovered from search string specifications, and number of usable papers.

| Term | Search String | Initial Papers | Usable Papers |
| --- | --- | --- | --- |
| Andromorph,  Gynomorph | TS=(andromorph* OR gynomorph*) NOT TS=(“human” OR “plant” OR “fungi” OR “bacteria”) NOT TS=(gynandromorph*) | 75 | 60 |
| Female Hormone,  Male Hormone | TS=(“female hormon*” OR “male hormon*”) NOT TS=(“human” OR “plant” OR “fungi” OR “bacteria”) | 47 | 40 |
| Female-like, Male-like | TS=(“female-like” OR “male-like”) NOT TS=(“human” OR “plant” OR “fungi” OR “bacteria”) | 124 | 124 |
| Feminize, Masculinize | TS=(“femin*” OR “masculin*”) NOT TS=(“human” OR “plant” OR “fungi” OR “bacteria”) | 580 | 539 |
| Gender / Sex Conflation | TS=("gender") NOT TS=("human" OR "plant" OR "fungi" OR "bacteria") NOT TS=("nomenclature" OR "women" OR "men" OR "sexuality" OR "gender bias" OR "society") | 1386 | 1047 |
| She-male, Transsexual, Transvestite | TS=(“transsexual*” OR “transvest*” OR “she-male”) NOT TS=(“human” OR “plant” OR “fungi” OR “bacteria”) | 10 | 7 |

Literature survey results

We initially identified 2,222 papers, 1817 of which were exclusive to non-human animals, could be assigned a taxonomic class, and were available to Loyola University Chicago online (Table S1).

Our data visualization reflects the raw data from our literature survey (Figure S1). We found a significant relationship between taxonomic classes and search terms in all searches except for feminize/masculinize and gender/sex conflation (Table S2). The terms andromorph/gynomorph were specific to Insecta, and especially the order Odonata. The use of ‘female-like’ and ‘male-like’ were most common in the class Aves (birds). Outdated terms such as “she-male” were specific to Reptilia (reptiles). For the terms feminize/masculinize, the classes Actinopterygii (bony fish) and Malacostraca (crustaceans) appeared more often than other taxes. The conflation of gender and sex was most common in the class Mammalia (mammals), followed by birds.

Table S2. Chi-square test of taxonomic bias for each search term from our literature survey.

| Term | χ2 | DF | P-value |
| --- | --- | --- | --- |
| Andromorph / Gynomorph | 160.15 | 15 | < 0.0001 |
| Female Hormone / Male Hormone | 41.08 | 15 | 0.00031 |
| Female-like / Male-like | 100.16 | 15 | < 0.0001 |
| Feminize / Masculinize | 22.32 | 15 | 0.0996 |
| Gender / Sex Conflation | 10.53 | 15 | 0.78 |
| Outdated Terms (transsexual, transvestite, she-male) | 172.9 | 15 | < 0.0001 |

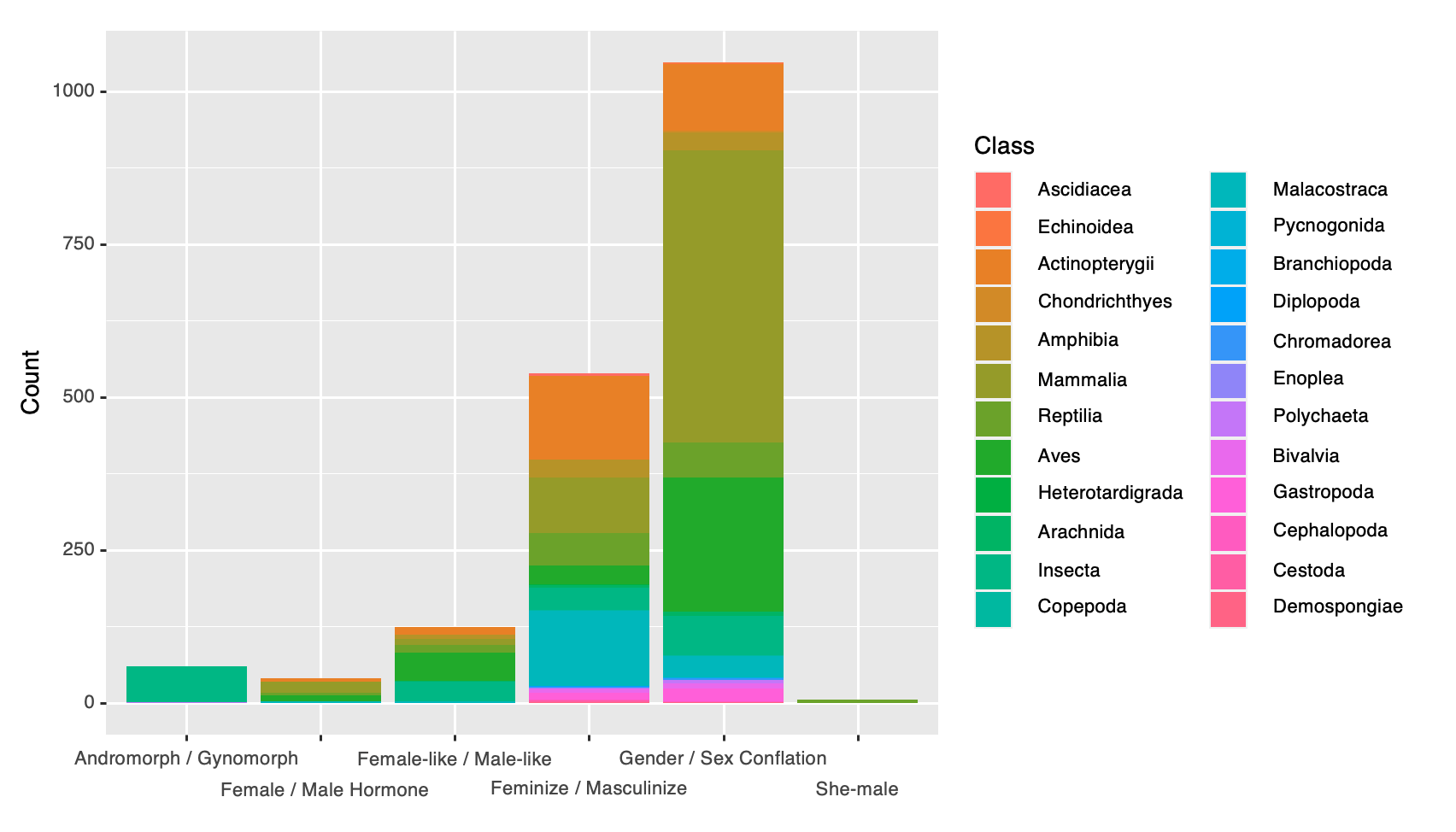

**Figure S1.** Count data from literature survey for terms organized by taxonomic class.

We found a significant increase in the presence of terminology over time, for all terms except “she-male” (Figure S2).

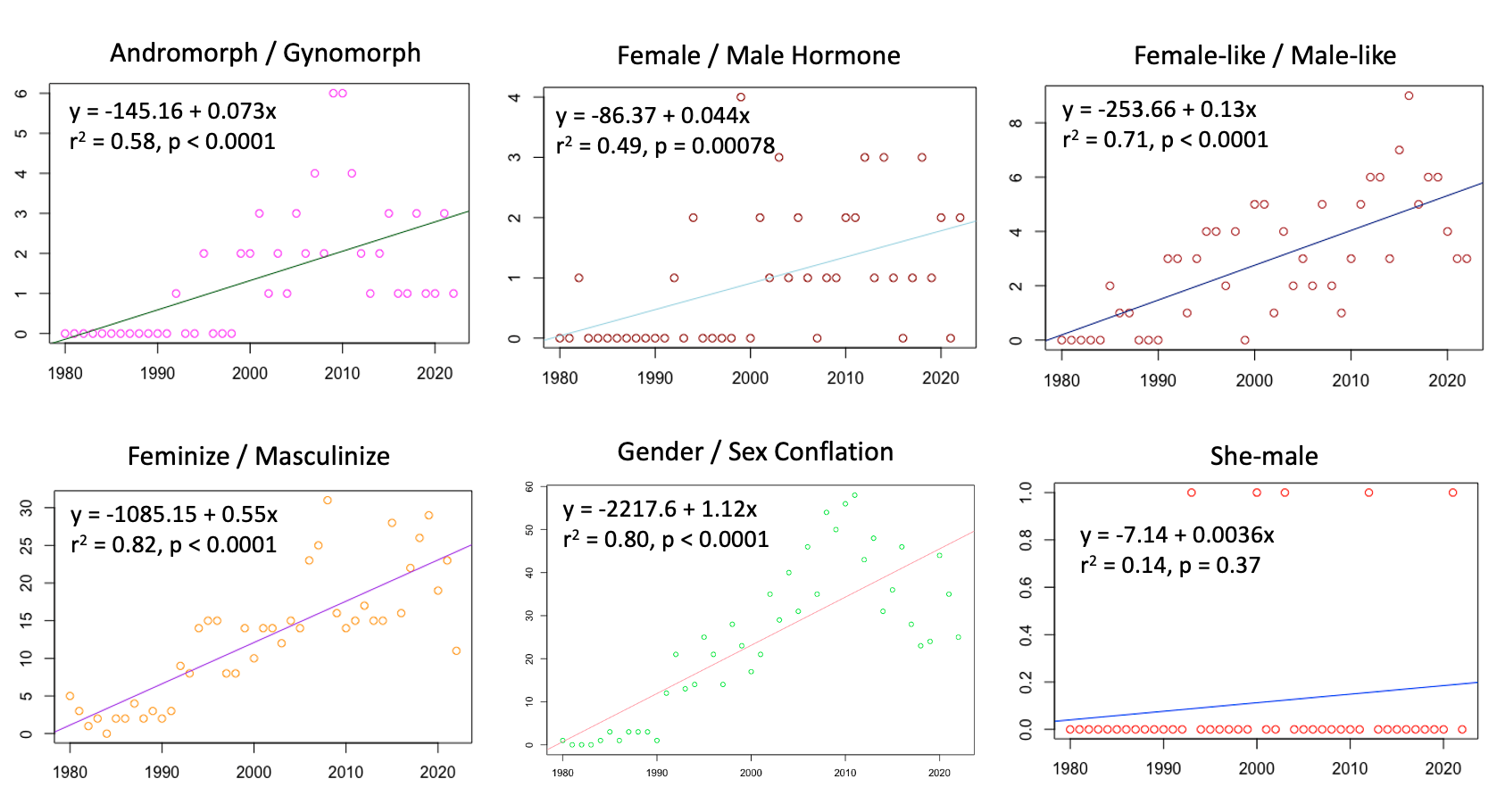
**Figure S2:** Terminology prevalence over time, with lines of best fit.
